# Using ultraviolet absorption spectroscopy to study nanoswitches based on non-canonical DNA structures

**DOI:** 10.1101/2021.08.10.455856

**Authors:** Blair McCarte, Owen T. Yeung, Alexander J. Speakman, Alistair Elfick, Katherine E. Dunn

**Affiliations:** School of Engineering, Institute for Bioengineering, University of Edinburgh, The King’s Buildings, Edinburgh, EH9 3DW, Scotland, UK

**Keywords:** Non-canonical DNA structures, DNA nanoswitches, ultraviolet absorption spectroscopy, G-quadruplexes, DNA triplexes

## Abstract

Non-canonical forms of DNA are attracting increasing interest for applications in nanotechnology. It is frequently convenient to characterize DNA molecules using a label-free approach such as ultraviolet absorption spectroscopy. In this paper we present the results of our investigation into the use of this technique to probe the folding of quadruplex and triplex nanoswitches. We confirmed that four G-quartets were necessary for folding at sub-mM concentrations of potassium and found that the wrong choice of sequence for the linker between G-tracts could dramatically disrupt folding, presumably due to the presence of kinetic traps in the folding landscape. In the case of the triplex nanoswitch we examined, we found that the UV spectrum showed a small change in absorbance when a triplex was formed. We anticipate that our results will be of interest to researchers seeking to design DNA nanoswitches based on quadruplexes and triplexes.

**Highlights:**

- Ultraviolet absorption spectroscopy can probe non-canonical DNA structures.
- Absorbance at 295nm tends to increase as G-quadruplexes form.
- Four G-quartets are needed to form a quadruplex with less than 1mM potassium.
- Formation of DNA triplexes can also yield a small change in UV spectra.
- UV absorption is a cheap label-free method for studying DNA nanoswitches.

## INTRODUCTION

The ability to manufacture customized DNA sequences underpins the field of DNA nanotechnology (1–4), which involves the use of synthetic DNA strands to make self-assembling nanostructures, devices and materials. DNA nanoswitches are small DNA molecules that can be switched from one configuration to another by a stimulus, such as the addition of a specific DNA strand (5, 6), a change in pH (7–9) or the binding of a ligand (10), among others. DNA nanoswitches can be used to selectively open and close drug delivery capsules (11–13) and they can also play a role in molecular computing systems (14) , which offer scope for low-power information processing in a biocompatible framework (15). In addition, DNA nanoswitches find applications in a range of biosensors (16, 17).

There is increasing interest in the use of non-canonical DNA structures for nanotechnology, including quadruplexes (18) or triplexes (19) or both (20). G-quadruplexes occur within or between DNA molecules that are G-rich. Four guanine bases can hydrogen bond to each other to form a G-tetrad, and a stack of such tetrads forms a quadruplex, which may involve 1, 2, 3 or 4 separate DNA strands (21). G-quadruplex formation is favoured by the presence of cations, particularly K^+^ (22, 23). There are several different topologies for G-quadruplexes, and the topology formed is influenced by the salt concentration (24).

Triplexes are formed from 3 DNA segments, and may involve 1, 2 or 3 separate strands. A triplex comprises a normal DNA double helix (with conventional AT/GC base-pairing), and an additional strand that occupies its major groove, binding to the duplex through Hoogsteen interactions (25) and forming nucleotide triplets. Triplexes can form with the additional third strand in either a parallel or antiparallel direction, and different triplet configurations are possible (26). In the present paper, we refer to parallel, major groove triplexes, which use a pyrimidine-rich third strand to form TAT or C^+^GC triplets. Due to the required protonation of cytosine to form these triplets, the formation of these triplexes is both sequence-specific and pH-dependent, and the transition pH can be tuned by changing the base content (9).

In biology, non-canonical DNA structures such as G-quadruplexes and triplexes have been implicated in gene regulation and may have therapeutic applications (27–30). It has been demonstrated very recently using single-molecule fluorescence microscopy *in vivo* that G-quadruplexes are associated with processes such as transcription and regulation, utilizing a fluorogenic quadruplex-binding ligand (31). Fluorescence measurements can also provide insights into the design of nucleic acid nanoswitches for sensing and computing applications. For example, transitions between the open and closed state can be probed using a modified nanoswitch in which a fluorophore is placed at one site in the nanoswitch and a quencher at another. The fluorophore and quencher must be selected and placed such that energy transfer is possible between them, but only when the switch is closed and they are brought into close proximity. Hence, the fluorescence emission is high when the switch is open and low when it is closed. The quencher can be replaced by a second fluorophore that is a FRET partner to the other, and the FRET signal measured. The general principle of this type of experiment is very well-established in the field of DNA nanotechnology and has been used in a multitude of studies (5, 9, 32, 33), including the investigation of nanoswitches based on non-canonical structures (7–9). In the case of G-quadruplexes, it is also possible to exploit the fact that G nucleobases tend to quench fluorescence of nearby dyes, and this phenomenon was used by Zhang *et al.*in the design of a potassium sensor (34). In the absence of potassium, their DNA construct took the form of a duplex. After addition of potassium, the formation of a G-quadruplex disrupted the duplex and released a strand carrying a fluorophore, which had previously been quenched by the guanines.

The major disadvantage of fluorescence is the need to add modifications such as fluorophores and quenchers. From a typical DNA synthesis company, a single-stranded unmodified 25nt DNA molecule would cost £12.00/$16.67/€14.03 at the 100 nmole synthesis scale [https://eu.idtdna.com/], but the addition of a 5’ terminal Cy5 would increase the cost to £95.00/$132.06/€111.11, even though Cy5 is one of the cheaper fluorophores. These prices are correct as of 2^nd^ August 2021 (prices generated in £, converted using exchange rate) include the extra expense of the HPLC purification required, and exclude any applicable taxes or institutional discounts. A dual-labelled strand (with Cy5 at 5’ end and a black hole quencher at the other) could cost £200.00/$277.90/€234.00. Consequently, there is a strong case for using label-free techniques.

Different G-quadruplexes display different features in circular dichroism (CD) spectra (35), which have been used to investigate the critical concentration of potassium required for folding, the presence of multiple G-quadruplex topologies in a single sample and the structural effect of ligand binding (36). However, DNA nanotechnology research groups often do not have convenient access to CD spectrometers, while they do usually possess UV-visible spectrometers, which have been used to probe non-canonical DNA structures. For instance, Mergny *et al.* performed UV absorption spectroscopy experiments in which they observed the transition between G-quadruplexes and a single-stranded random chain conformation as DNA samples were heated or cooled (37), using sequences observed in biological systems (mainly telomeric repeat sequences). They demonstrated that UV-absorbance spectra of G-quadruplexes have a characteristic signature at 295nm and that the melting temperature of the structure was significantly affected by the nature of the monovalent ion present in the buffer. Their results showed that potassium contributed most to the stability of quadruplexes, and sodium had a weaker stabilizing effect, with lithium being even less effective than sodium. Later, Karsisiotis *et al.* investigated quadruplex technology using a similar technique in combination with circular dichroism (CD) (38). They obtained thermal difference spectra and measured the differences in absorbance at specified wavelengths (*λ*) at high temperature (melted) and low temperature (folded). Their results showed that ΔA_240_/ΔA_295_ was considerably higher for parallel than antiparallel quadruplexes, and that further information on topology could not be obtained from UV absorption measurements. Zhang & Balasubramanian used UV absorption measurements to study four specific quadruplex forming oligonucleotides (three of RNA, one of DNA), and obtained titration curves showing absorption changes as a function of potassium ion concentration (39). It has recently been demonstrated that G-quadruplexes with different structures/topologies can be distinguished by performing a competition assay, in which dye molecules can bind either to the quadruplexes or to a range of ‘hosts’, with the fluorescence signal indicating the type of quadruplex present (40).

DNA triplexes have also been studied using CD and UV absorption spectroscopy. For instance, CD has been used to show that triplex formation can be detected in comparison to the spectra for the original sequence when no triplexes are formed. However, no characteristic triplex spectral features have been identified conclusively, meaning that CD has to be performed for each sequence individually in both triplex and non-triplex forming conditions to indicate triplex formation. It may also be necessary to acquire data using a complementary technique such as gel electrophoresis (41). Brucale *et al* studied a set of triplex forming DNA oligonucletotides and changed the pH of the solution to shift between triplex-forming and non-triplex-forming conditions. Under low pH (triplex-forming) conditions they observed a small drop in absorbance at 260 nm(42) .

In this paper, we describe the results of our investigation into the use of UV spectroscopy to characterize the switching of G-quadruplex and triplex DNA nanoswitches at fixed temperature, driven by salt concentration and pH respectively. Our first objective was to extend the previous studies and further validate the use of this technique for studying such nanoswitches, and our second objective was to ascertain how the sequence used in the nanoswitch affected the conditions under which it would switch. As is common in the field of DNA nanotechnology, we used designed sequences rather than those of biological origin.

As expected, we found that the absorbance at a wavelength of 295nm changed systematically with increasing salt concentration for the strands designed to form G-quadruplexes. Results obtained by two independent experimentalists were consistent. Control experiments showed that the absorbance at 295nm did not depend on salt concentration for a reference strand that contained only thymine nucleotides. For sequences consisting of G-tracts separated by spacers consisting of three thymines, we found that the strands with longer G-tracts required fewer cations to stabilize them. We also observed that Na^+^ ions enhanced the stability less than K^+^ ions, in accordance with the previous studies mentioned above. We also found that changes in the spacer region could completely disrupt G-quadruplex formation, and this is attributable to traditional base-pairing interactions. This result underlines the importance of eliminating interactions that could compete effectively with the quadruplex-forming process.

For triplex nanoswitches, we found that the UV spectra showed a drop in absorbance at both 240 nm and 260 nm when a 10-nucleotide long triplex was formed, which indicates that for some triplex-forming constructs the UV spectra can indeed be used to study the transition between structures. This method of detection may prove useful when other more established approaches such as using fluorophore labelled oligonucleotides or circular dischroism spectra are not appropriate or are inaccessible due to equipment or budget limitations.

## METHODS

### DNA sequences: G-quadruplexes & control

We used five DNA strands, four of which were designed to form G-quadruplexes, based on previous studies (34, 37). As shown in Fig. 1 and Table 1, the strands differed in the number of G-quartets that could form or the sequence of the spacers between G-tracts. The fifth strand was a 28-nucleotide poly-thymine strand, which served as a control, to confirm that observed effects were attributable to quadruplex formation.

**Figure 1.**
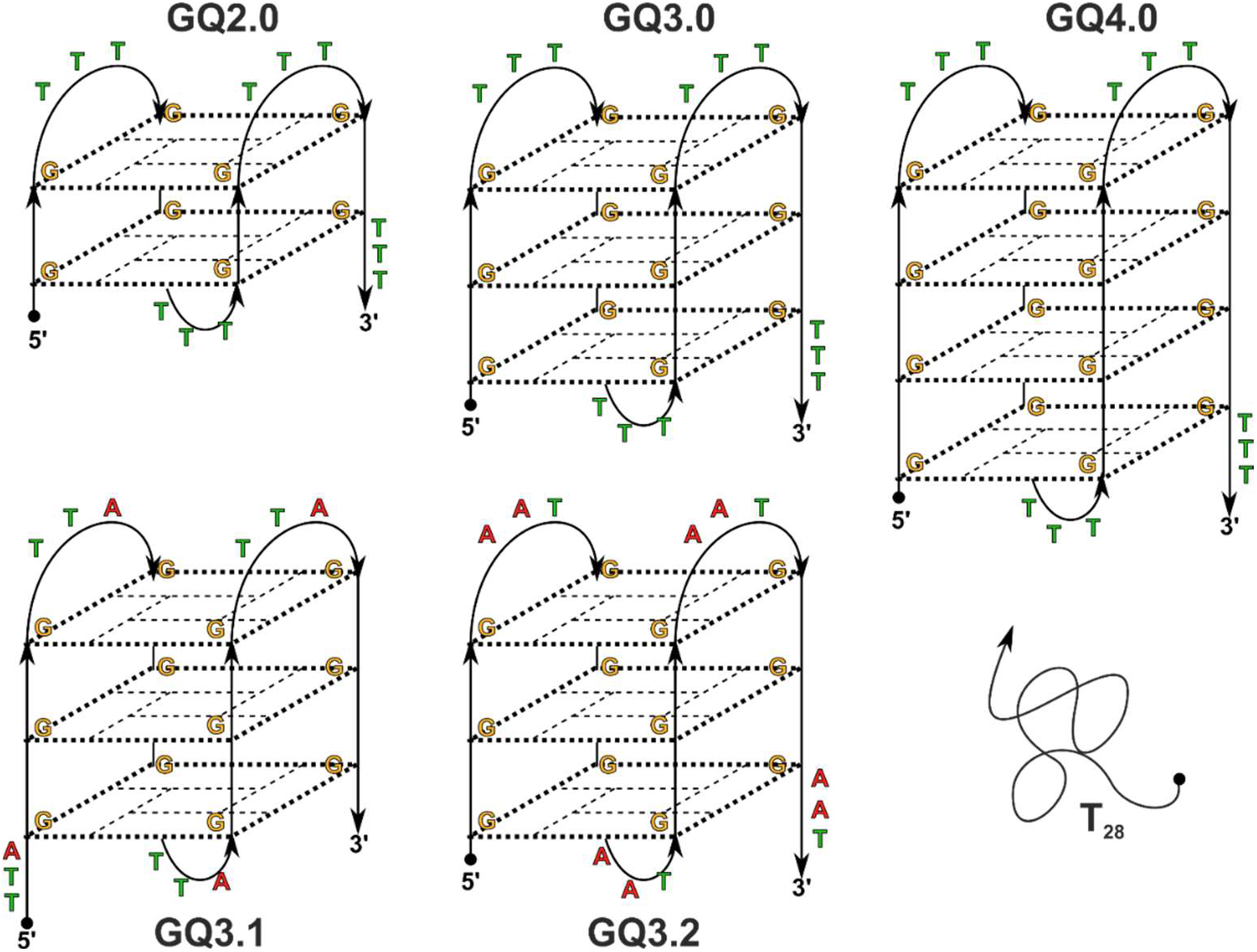
Schematic diagrams of G-quadruplex structures, showing variations in loop sequences and number of G-quartets. Images do not necessarily represent the correct folding topology; antiparallel basket topology is assumed for illustrative purposes.

**Table 1:**
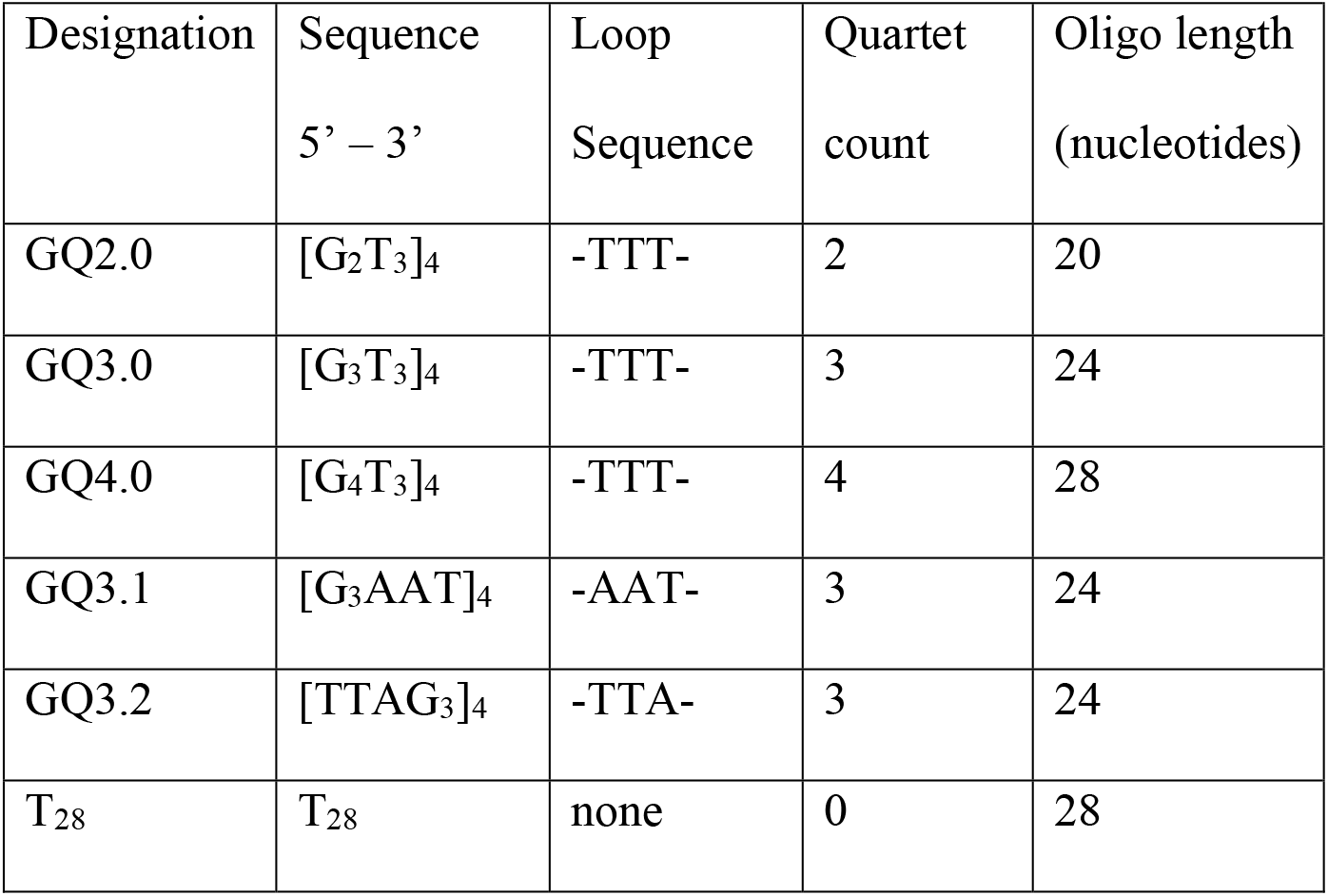
Sequences of DNA oligonucleotides used for the G-quadruplex experiments, including the polythymine control.

Oligonucleotides (Table 1) were purchased from Integrated DNA Technologies (IDT) with no additional purification. Lyophilized DNA was resuspended in 1xTE buffer to a final concentration of 100 μM. Resuspended DNA was stored at −20 °C.

### UV absorption spectroscopy: G-quadruplexes & control

All work was carried out at room temperature. UV absorption measurements were taken using a Nanodrop 2000c spectrophotometer, operated in pedestal mode. The ‘path length’ in the software was set to 1mm. The pedestal sensor was cleaned before each reading as follows. 2μl of Type 1 (ultrapure) water (UPW) was pipetted onto the sensor and the measurement arm was lowered. The measurement arm was then returned to a raised position and liquid removed from both sides using Kimtech Delicate Task wipes. The cleaning step was performed twice before 2μl of reference buffer (1xTE) was loaded onto the measurement pedestal and the blank measurement was taken. The sample was removed using Kimwipes before 2μl of the DNA sample was loaded and measured. Repeat measurements were taken consecutively, with each sample being removed using Kimwipes.

The linear range of the spectrophotometer was determined using samples of GQ3.0 ([G_3_T_3_]_4_). DNA concentration standards from 2.5-100 μM were prepared by dilution of a 100 μM GQ3.0 stock solution in 1xTE (with no salt). A 1.25 μM sample was prepared by dilution of the 2.5 μM sample in 1xTE. For each concentration analysed a total of five measurements were taken.

Two individuals acquired UV spectra for G-quadruplex and control samples at different salt concentrations. The original (preliminary) dataset focussed on G-quadruplexes GQ3.2 (with KCl or NaCl), GQ4.0 (with KCl), GQ2.0 (with KCl). The second, more complete dataset comprised the results from comprehensive tests of all five sequences for both salts (including the poly-thymine control). The results presented in this paper are taken exclusively from the second dataset.

G-quadruplex nanoswitch folding was induced by exposure to sodium/potassium ions in solution. Salt stock solutions were prepared at concentrations of 10mM, 100mM and 500mM for both NaCl and KCl in 1xTE buffer. Salts were purchased from Sigma Aldrich. 50μl folding samples were prepared in 1.5ml Eppendorf tubes with different salt concentrations. The DNA concentration was the same for all samples (20 μM), and the salt concentration varied from 0.2 mM to 100 mM (NaCl) or 1 mM to 250 mM (KCl). Blank samples were produced using similar recipes in which the 10 μl of 100 μM stock DNA solution was replaced with 1xTE buffer. DNA was diluted in 1xTE before the salt was added.

For each G-quadruplex-forming experiment at least six individual measurements were performed. Further repeats were carried out if anomalies were observed after data processing.

### DNA sequences: triplex experiments

The triplex-forming molecule is shown in Fig. 2. At high pH the construct consists of a 19bp duplex with a 56nt single-stranded tail. At low pH the final 10nt segment of the tail binds within the major groove of the terminal 10bp of the duplex, forming a triplex. The singlestranded spacer comprised a 25nt long linker region designed using EGNAS (43) to avoid forming secondary structures. Our triplex-forming molecule was based on that used by Idili *et al.* (9), but we added extra nucleotides in both the double stranded and single stranded spacer region. This modification shifted the transition midpoint from approximately pH8.5 (9) to 6.3, according to our fluorimetry experiments. The constituent oligonucleotides of the triplex-forming construct were purchased from Integrated DNA Technologies (IDT) with no additional purification. Upon receipt, oligonucleotides were resuspended in ultrapure water to a concentration of 100 μM.

**Figure 2.**
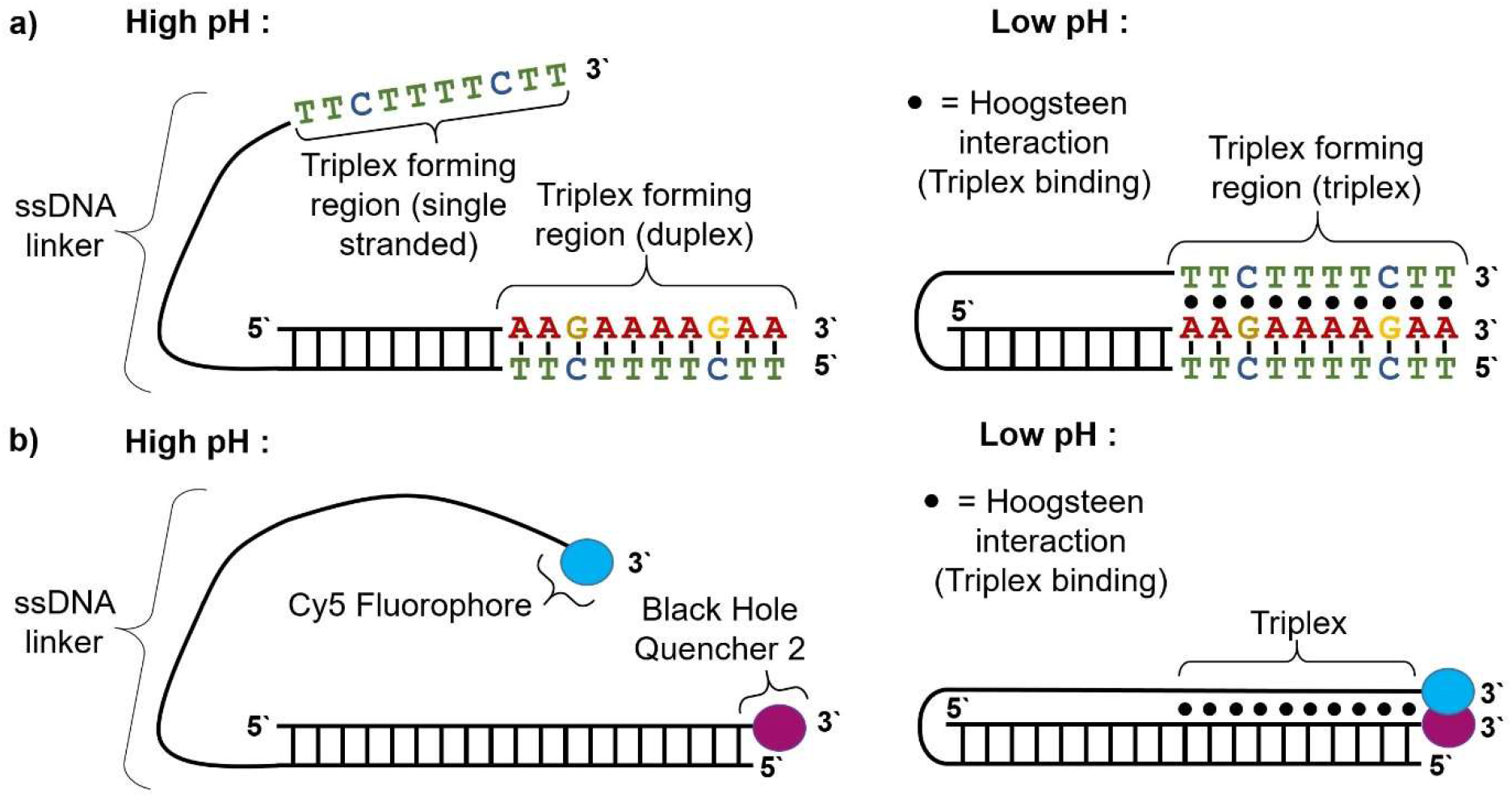
Triplex nanoswitches used in this investigation. a) Schematic diagrams of Triplex structures, showing their formation via Hoogsteen interactions at low pH. b) Mechanism of triplex detection in fluorimetry experiments, illustrating how a fluorophore (Cy5) can be brought into proximity of a quencher (BHQ2) when the triplex forms, resulting in a drop in fluorescence.

### Britton Robinson buffers: triplex experiments

Britton Robinson buffers were produced at 11 different pH levels from 5 to 11.74 by mixing set volumes of 0.4M hydrochloric acid, 0.4M boric acid (ACROS Organics, #327132500), 0.4M acetic acid (ACROS Organics, #10005920), 0.4M phosphoric acid (Alfa Aesar, #11459216), 0.04M MgCl2 (ACROS Organics, #197430010) with variable volumes of sodium hydroxide at either 0.2M or 0.4M (Fisher Scientific, #10745141) and ultrapure water, according to the recipe provided by Cerdà and Mongay (44) as specified in the Supplementary Information. pH was measured using a Sartorius pH meter (model #PP-15).

### Sample preparation and UV absorption spectroscopy: triplex experiments

100μM stock oligonucleotides were mixed in a 1:1 ratio to form a solution of 50μM of both oligonucleotides. 2μL of this stock was added to 18μL of Britton Robinson buffer of appropriate pH, to produce a final DNA concentration of 5μM. 2μL of ultrapure water was added to the same buffer for use as a blank for absorption measurements. Samples were placed on a slow-moving shaker and left to hybridize for 10 minutes before being analysed by UV absorption spectroscopy, which was performed with 6 replicates for each using the same method as for G-quadruplexes.

### Fluorescence spectrophotometry: triplex experiments

To confirm triplex formation and ascertain the pH midpoint of the transition, appropriate sequences were ordered from IDT with fluorophore and quencher labels as shown in Fig. 2 and Table 2. As a control, a non-triplex forming oligonucleotide was purchased with an alternative 3’ terminal 10nt sequence (Table 2), designed using EGNAS (41) to avoid forming secondary structure in the linker region. Oligos were suspended in ultrapure water to a concentration of 100μM. 0.5μL of long Cy5-labelled oligo (either triplex forming or control) and 0.5μL of short, BHQ2-labelled oligo was added to 2mL of Britton Robinson buffer of a known pH and inverted repeatedly to mix. This was left for 5 minutes to hybridize, resulting in a final hybridized DNA concentration of 25nM. 2mL of this solution was then loaded into a quartz cuvette (Agilent, #6610000900) and scanned using a Cary Eclipse Fluorescence Spectrophotometer (Agilent, #G9800A). Settings used for the scan were as follows: excitation wavelength = 648nm, emission range = 660nm to 700nm, excitation and emission slit widths = 5nm, averaging time = 1s, PMT voltage = 800V. Scans were performed for both triplex-forming and control samples at 23 different pH levels with 3 replicates for each. The peak fluorescence emission was observed at 664nm, and the fluorescence at this wavelength was isolated from each scan and used for later comparison.

**Table 2:**
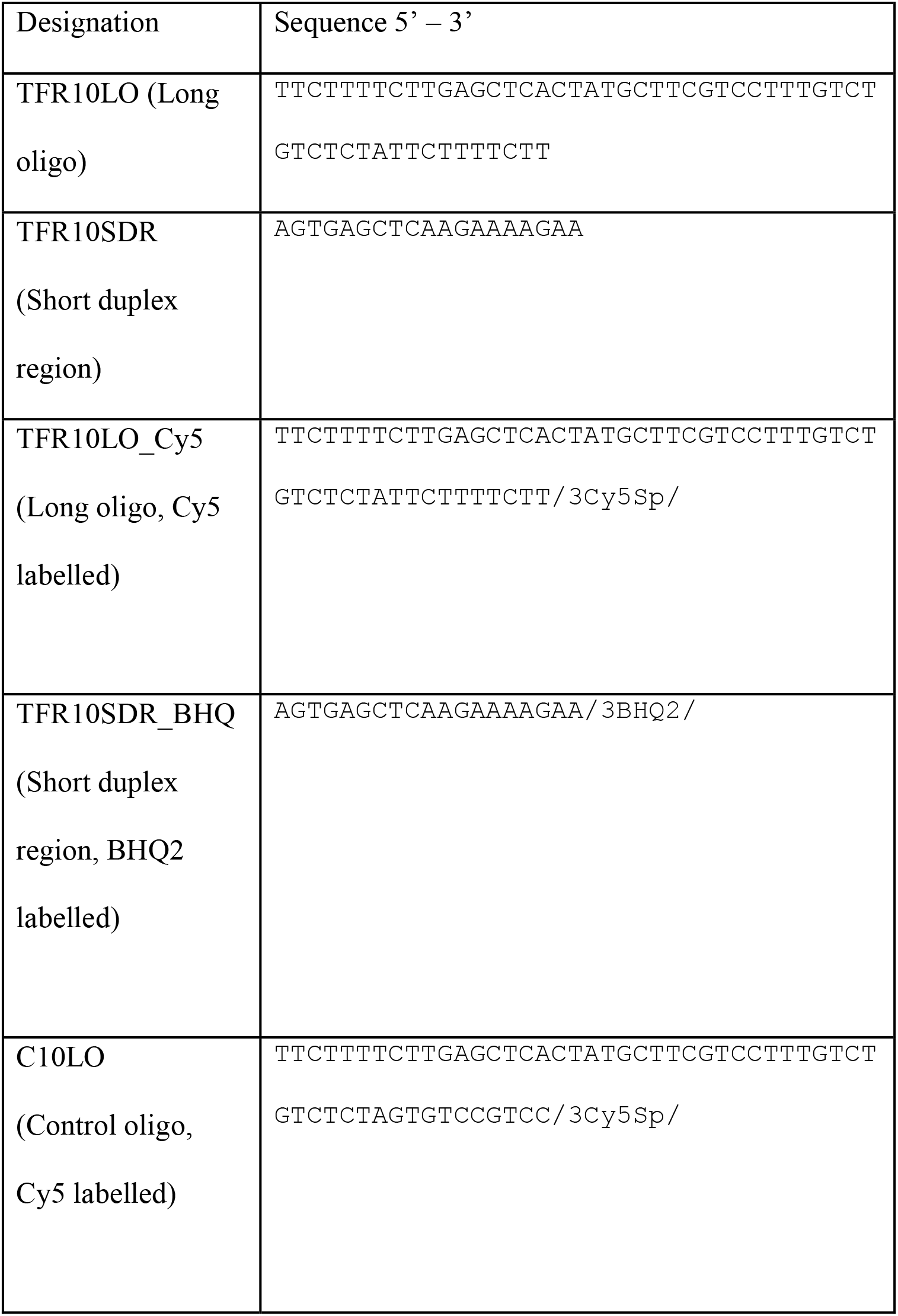
Sequences of DNA oligonucleotides used for the triplex experiments.

### Data analysis

The preliminary G-quadruplex dataset was processed manually whereas the second dataset was processed automatically using Python 3.7.3 and MATLAB and scripts are available on github. For the triplex dataset, data was processed manually. All data and scripts used for this paper are available in the supplementary files.

## RESULTS & DISCUSSION

### Spectrophotometer Linearity

We determined suitable working DNA concentrations by ascertaining the range of concentrations for which absorbance at 260nm depended linearly on concentration, using the strand GQ3.0 [G_3_T_3_]_4_ (Fig. 1). We used our spectra (Fig. 3a) to establish the region in which the spectrometer response was linear (Fig. 3b). At higher concentrations the response started to deviate from linear behaviour, as expected. For 80*μ*M and 100*μ*M, increasing distortion was also visible around the 260 nm peak (Fig. 3a), suggesting detector saturation and potential issues with automatic correction for low transmittance. Slight deformation could be seen in the 40*μ*M curve as the absorbance approached 1. On the basis of these results, we chose 20*μ*M as a suitable maximum DNA concentration.

**Figure 3:**
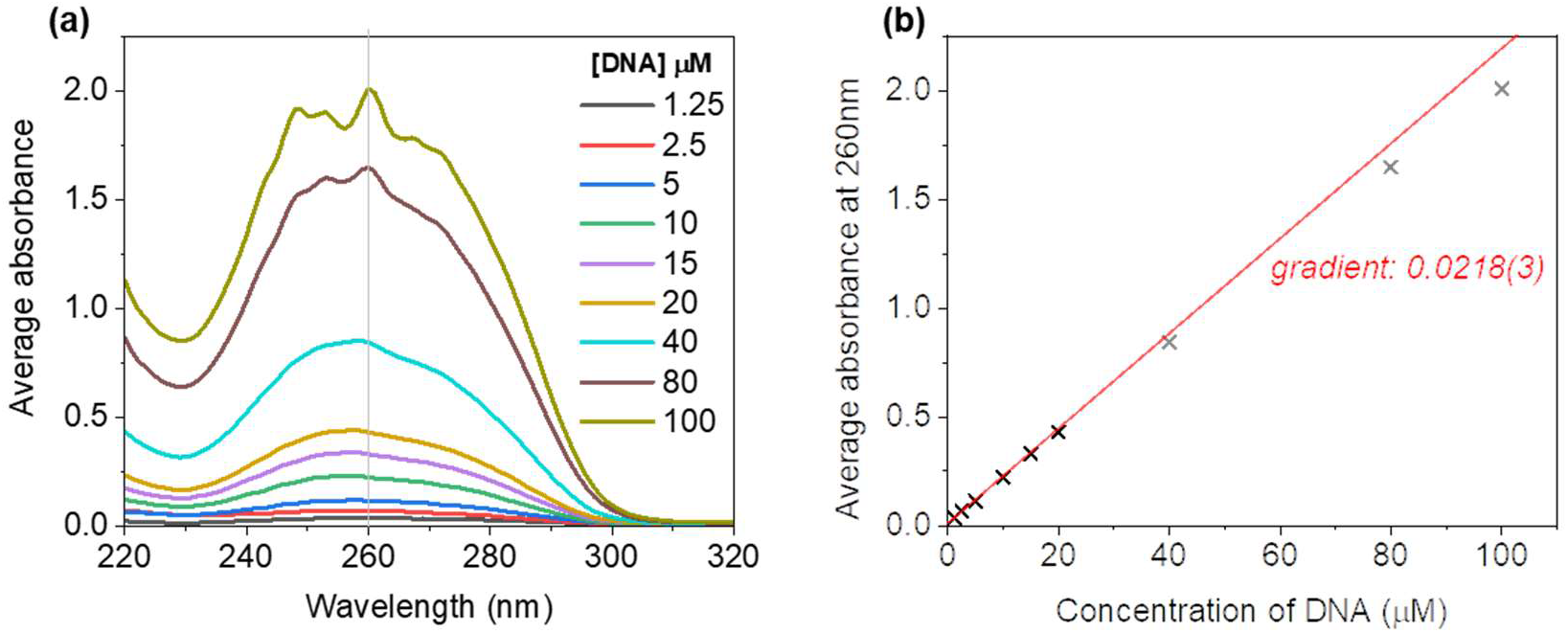
Spectrophotometer linearity & optimal DNA concentration. (a) Spectra (220-320nm) for samples of the DNA sequence GQ3.0 ([G_3_T_3_]_4_) at concentrations of 1.25μM to 100μM in 1xTE. The displayed data is an average of 5 measurements and path length was 1mm. (b) Extracted average absorbance at 260nm as a function of DNA concentration. Red line: linear fit, with intercept fixed at zero. The three grey data points were omitted from the fit.

Anecdotally, some laboratories regard Nanodrop measurements as unreliable. Our experience shows that this is incorrect, as long as the pedestal is cleaned thoroughly between runs (Methods), repeat measurements are taken, and the instrument is operated in the range for which the response is linear. For validation purposes, we compared our results to the predictions of basic theory. According to Beer’s law, the absorbance, *A*, is equal to the product of the extinction coefficient, *e*, the concentration, *c*, and the path length, *b*. For these measurements, the effective path length was set in the software to be 0.1cm. According to an online calculator provided by our DNA supplier (https://eu.idtdna.com/calc/analyzer), the extinction coefficient for the oligonucleotide GQ3.0 was 222000 L/(mole·cm) = 0.222μM^−1^cm^−1^, giving a predicted value for the gradient (=eb) of 0.0222 μM^−1^, which is almost identical to the experimentally obtained value of 0.0218(3) μM^−1^.

### G-quartet Induced Absorbance Shift at 295 nm

Fig. 4a shows the absorbance spectra of GQ3.2 ([TTAG_3_]_4_), for low (0.2mM), medium (2mM) and high KCl concentrations (100 mM). To produce each spectrum, six repeat measurements were averaged, prior to normalization to the value at 260nm. A clear increase in absorbance at 295 nm was observed as KCl concentration was increased, and we attributed this to G-quadruplex formation, which is favoured by the presence of potassium ions.

**Figure 4:**
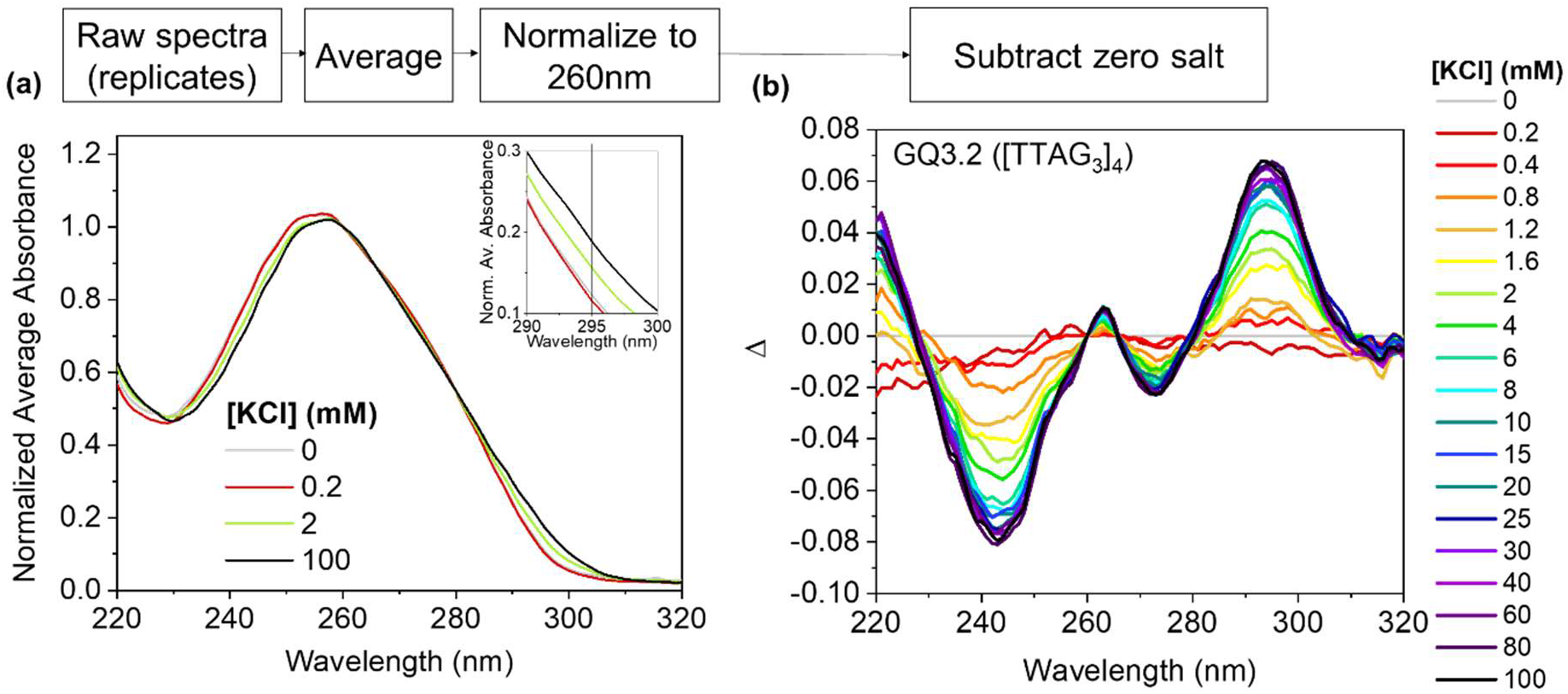
Spectra for GQ3.2 for different concentrations of KCl, with fixed concentration of DNA. (a) Absorbance spectra (averaged over 6 datasets, then normalized to the value at 260nm). Inset shows close-up of the region around 295nm. Absorbance is dimensionless. (b) Difference spectra for GQ3.2, produced by subtracting the normalized average spectrum for the zero salt case from the corresponding spectrum for each concentration of salt. We define Δ = normalized(average absorbance) _salt_ − normalized(average absorbance)_zero salt_

The changes were more conspicuous in the ‘difference spectra’ (Fig. 4b), computed by subtracting the average spectrum for the zero salt case from the corresponding data for each concentration of KCl. Prior to the subtraction, the average spectra were normalized to the 260 nm absorbance maximum. The difference spectra had several peaks that appeared to strengthen as the potassium concentration was increased. In line with the previous studies mentioned above, we focused on the peak at 295nm. The difference spectra also featured isosbestic points (where the absorbance does not depend on the salt concentration), such as that at 280nm, noted in studies by Mergny *et al* (37)and Scaria *et al* (45).

In our difference spectra, we saw almost no signal for KCl concentrations of 0.2mM and 0.4mM, which indicated that very little folding occurred at such low levels of potassium for this strand. As we increased the concentration from 0.2 mM to 6 mM, we observed a dramatic increase in the difference of the normalised absorbance at 295nm. Above 10mM of K^+^, very little further increase was apparent.

### Effect of changing the number of G-quartets

We studied the effect of changing the number of G-quartets using three different G-quadruplex-forming strands, in both KCl and NaCl (Fig. 5a/b). As a control, we also performed the experiments with a 28-nucleotide poly-thymine oligonucleotide (T_28_), which is incapable of forming G-quartet structures or other secondary structure. As expected, the control measurements showed no change in absorbance at 295 nm in the presence of either sodium or potassium (Fig 5a/b).

**Figure 5:**
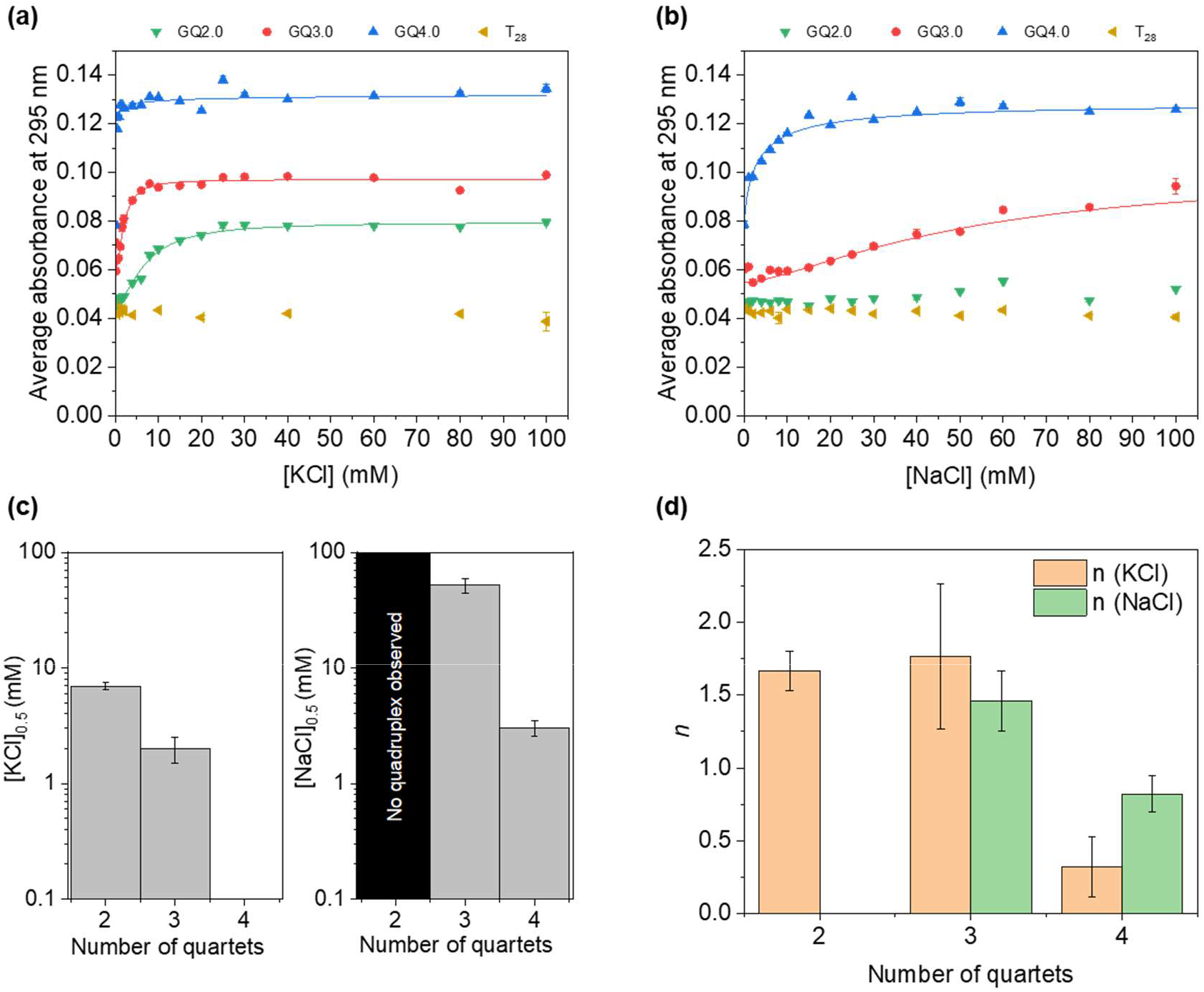
Salt-dependent G-quadruplex formation with different numbers of G-quartets. (a,b) Absorbance at 295 nm (averaged over 6 datasets) for strands that form G-quadruplexes with 2, 3 or 4 quartets (GQ2.0, GQ3.0 and GQ4.0 respectively). Absorbance is plotted as a function of the concentration of KCl or NaCl in the sample buffer. Control measurements for a poly-T strand (T_28_) are also shown. In most cases error bars are too small to be seen. Solid lines: fits performed with Origin using the equation <A_295_> = (START) + (END-START) x^n^) / (k^n^ +x^n^).The parameter ‘START’ was fixed. x represents the concentration of salt (either KCl or NaCl). (c) Salt concentration at the transition mid-point, equal to k^n^ Value and error bars were obtained using the results of the fit parameters. No quadruplex was observed for the NaCl – 2 quartet case, and thus no fit was performed in this case. (d) The value of the coefficient n for the five fits.

GQ4.0, GQ3.0 and GQ2.0 are designed to form quadruplex structures with 4, 3 and 2 G-quartets, respectively. In almost all cases, increasing the concentration of ions caused the average absorbance at 295nm (<A_295_>) to increase, as the quadruplexes folded (Fig. 3a/b). As expected, the value of <A_295_> was correlated with the number of G-quartets that could form. At high concentrations of ions, <A_295_> was greatest for GQ4.0 (4 quartets), followed by GQ3.0 (3 quartets) then by GQ2.0 (2 quartets).

We observed that the amount of potassium needed to induce folding was also dependent on the number of quartets – the fewer quartets, the more potassium required to stabilize the quadruplex. For GQ4.0, folding occurred at very low concentrations of potassium. A similar pattern was apparent for sodium (Fig. 5b). The data suggested that folding of the two-quartet quadruplex was incomplete in NaCl, because <A_295_> hardly changed over the entire range of [NaCl], and its value was significantly lower than that obtained in KCl. In all cases, K^+^ was more effective than Na+ for stabilization of G-quadruplexes.

Zhang & Balasubramanian (39) noted that a Hill function can be used to describe how <A_295_> depends on the concentration of potassium, and we used the same model. We therefore fitted our data with the following equation:

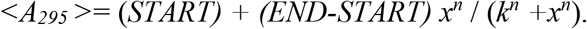

Here, *x* represents the concentration of salt (either KCl or NaCl), while *k*, *n*, *END* and *START* are fit parameters. The quantity *k^n^* corresponds to the concentration of salt at which half the quadruplexes were folded. Technically, this model implies an assumption of two-state folding kinetics, which is likely to be an oversimplification. However, as shown in Fig. 5a/b, the Hill function was an excellent fit for all cases in which folding was observed. This suggests that it is empirically useful even though the physics of the folding process is actually more complicated. For reference, a simple derivation of the two-state folding equation is provided in the Supplementary Information. The other assumptions are that the ions were present in excess, and all measurements were made at equilibrium.

Our fit results (Fig. 5c) indicated that 50% of GQ4.0 strands would be folded at a KCl concentration lower than the lowest non-zero KCl concentration for which we took a measurement, and the KCl transition midpoint is therefore effectively indistinguishable from zero. For NaCl, the transition midpoint did not occur until a salt concentration of 3±0.5mM was reached. For the quadruplex with three quartets (GQ3.0), the coefficient *n* was similar in both KCl and NaCl, indicating that the physics of the folding process was similar for both species of ion. In the case of GQ4.0, the coefficient for the KCl data took a value of only 0.32 but this is unreliable because the transition midpoint concentration was not significantly different from zero. For GQ2.0, no transition was observed in NaCl-based buffer. For KCl, the transition midpoint occurred at a concentration of 7.0±0.4mM, and the coefficient *n* was similar in value to that obtained for GQ3.0.

### Effect of changing sequence and position of the spacer

Each of the quadruplex-forming strands contained spacer domains that separate the G-tracts. When the G-quadruplex was folded, the spacer domains formed loops at the top or bottom of the quadruplex.

GQ3.0 and GQ3.1 both form quadruplexes with three quartets, and have the following sequence:

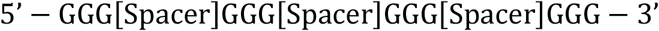

The spacer sequences were [TTT] and [AAT] for GQ3.0 and GQ3.1 respectively. In the case of GQ3.1, <A_295_> did not change significantly as the concentration of salt was increased, remaining at approximately the same value as seen for GQ3.0 in the absence of salt. Consequently, we deduced that GQ3.1 did not form a G-quadruplex structure at any of the salt concentrations we examined. We suspected that this might be due to the transient formation of weak secondary structure via conventional base-pairing, and used the online structure prediction tool NUPACK (46) to investigate this hypothesis. NUPACK predicted that GQ3.1 could potentially form a weak hairpin (Fig. 6c). The free energy of this secondary structure was very small (−0.55 kcal/mol), indicating that it would not be stable at room temperature. However, transient interactions involving these base pairs might be responsible for the disruption to G-quadruplex formation.

**Figure 6.**
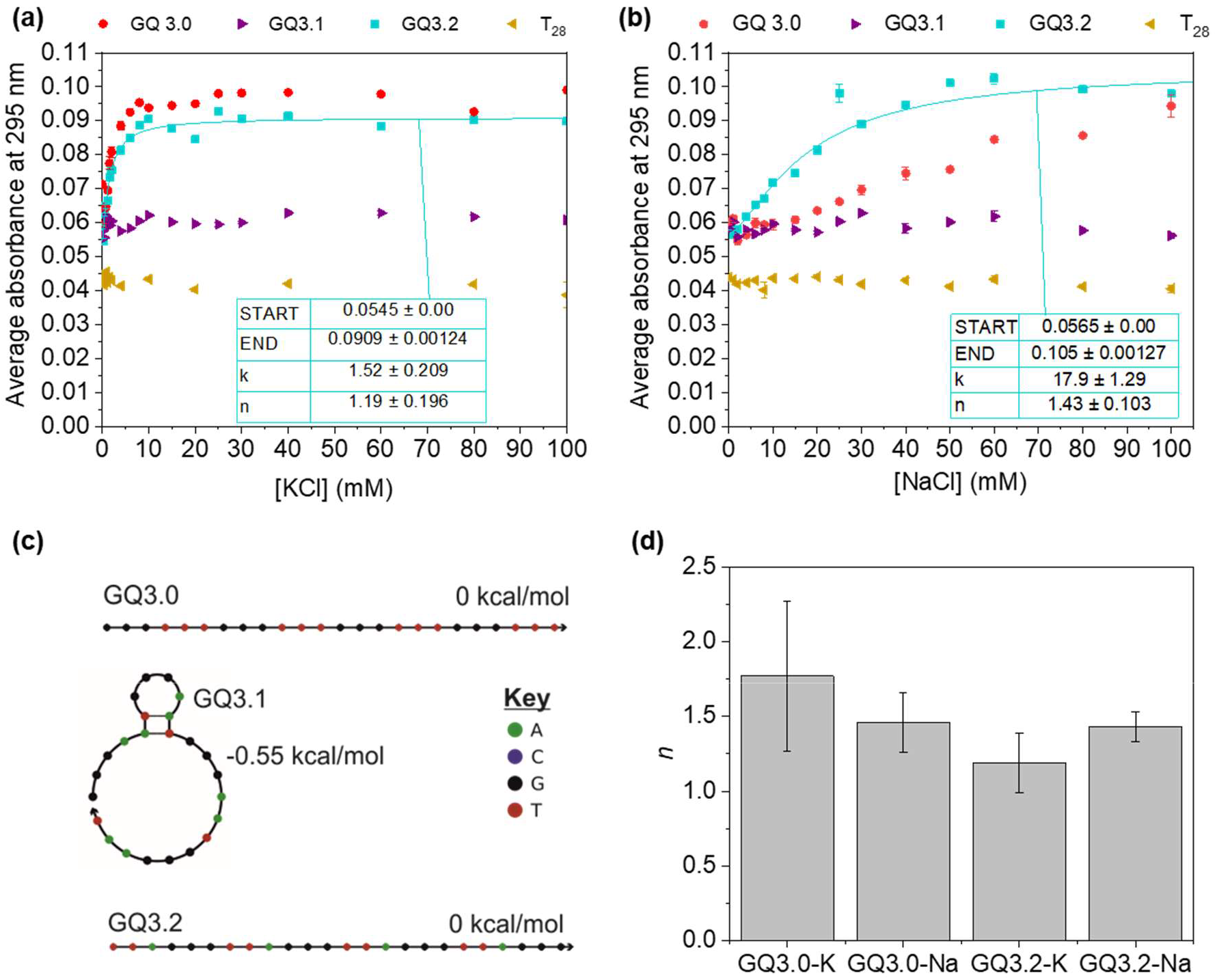
Salt-dependent G-quadruplex formation with three quartets and different spacers. (a,b) Absorbance at 295 nm (averaged over 6 datasets) for strands GQ3.0, GQ3.1 and GQ3.2. Control data for T_28_ is provided for reference. Formatting is consistent with that of Fig. 5a/b. Hill function fit is shown. (c) NUPACK predictions for secondary structure of the three strands of interest at 20°C for 1M Na^+^. (d) The value of the coefficient n for the indicated fits.

The strand GQ3.2 has the sequence [TTAGGG]_4_. This is almost the same as the hTelo sequence studied by Zhang & Balasubramanian (39), the only difference being that we added an extra ‘TTA’ at the 5’ end of the strand. Our value for the coefficient *n* for GQ3.2 (1.19±0.20, Fig. 6d) is consistent with the value they obtained for hTelo (0.9±0.1). However, our fit suggested that the folding transition midpoint occurred at 1.5±2mM KCl, whereas in their study hTelo was seen to fold at much lower concentrations of potassium. They performed the experiment by titration and corrected for dilution effects, whereas we prepared different samples for every salt concentration, with the same DNA concentration each time. We used a higher concentration of DNA, used pH8 buffer instead of pH7, and our oligonucleotide had an extra spacer (as noted above).

Our results showed that GQ3.0 and GQ3.2 underwent a folding transition at a similar concentration of KCl, but GQ3.0 required significantly more NaCl to fold than did GQ3.2. This may be attributable to different folding topologies.

### UV absorption spectroscopy measurements of triplexes

As described in the Methods section, we used a triplex construct based on that designed by Idili *et al.*, but we included additional nucleotides in the double and single stranded linker region, which was expected to lead to a decrease in the transition pH. We performed experiments using fluorophore and quencher labelled oligonucleotides, and we confirmed that a triplex was formed by TFR10 sequences (Fig. 7a) in a pH-dependent manner, while the C10 non-triplex control showed no signs of triplex formation (Fig. 7b). Using the emission fluorescence of C10 as a reference, we calculated the percentage of constructs in a triplex configuration at each pH level, giving an approximate triplex transition midpoint (where 50% of oligos have formed triplexes) of pH 6.3 (Fig. 7c). As expected, the transition occurs at a lower pH than observed by Idili *et al.* for their construct.

**Figure 7.**
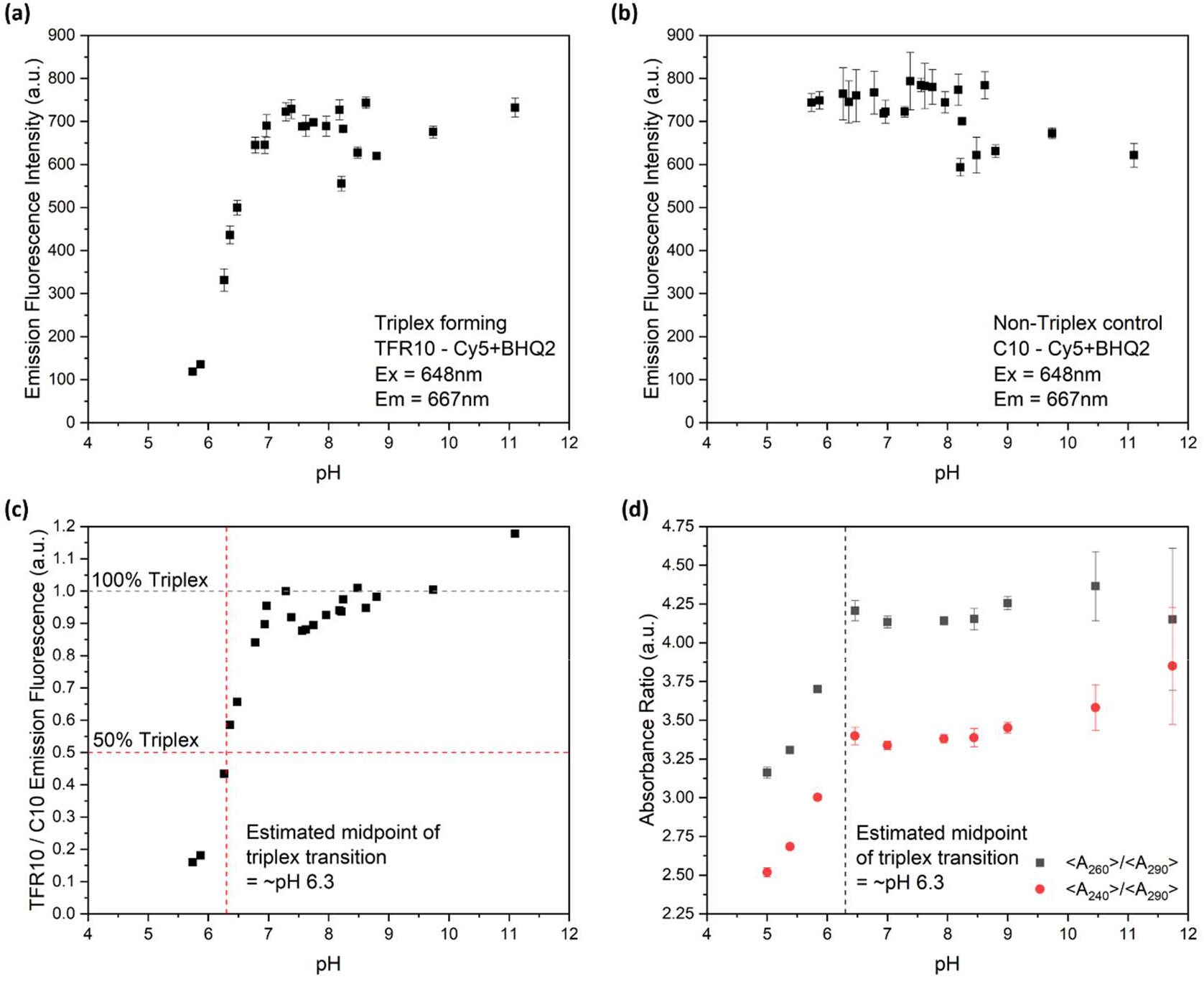
Fluorescence spectrophotometry and UV-visible absorbance measurements (averaged over replicates) of triplex-forming constructs and controls. (a) Fluorescence measurements made in different pH buffers on the TFR10 construct, which forms a 10nt triplex. TFR10 was labelled with Cy5 and BHQ2 such that fluorescence was suppressed under triplex-forming conditions. (b) Fluorescence spectrophotometry of C10, equivalent to TFR10 except for the substitution of a non-triplex forming sequence at the end of the singlestranded linker. (c) The fluorescence signal from the TFR10 construct divided by that obtained from C10. The transition midpoint is presumed to occur when this ratio takes a value of 0.5, estimated to be at approximately pH6.3. (d) UV absorption measurements at a range of pH levels, showing a drop in the ratio of absorbance values below the expected triplex transition midpoint, for both <A_260_>/<A_290_> and <A_240_>/<A_290_>, where the subscript denotes the wavelength in nm and <> indicates the average. Fluorescence measurements were taken in triplicate, while UV-visible absorbance measurements had 5 replicates each.

UV absorbance spectra of the same oligos (without fluorophore or quencher molecules attached) reveal a drop in both <A_240_>/<A_290_> and <A_260_>/<A_290_>. This occurs as the pH drops, most notably below pH 6.5 (Fig. 7d), which coincides with the transition observed in our fluorescence experiments. Our findings therefore indicate that triplex formation is associated with a drop in absorbance at 240nm and 260nm, compared to 290nm.

## CONCLUSIONS

In previous studies it had been shown that G-quadruplex structures produce characteristic features in UV spectra, giving rise to increased absorbance at around 295 nm. In this paper we extended those studies by accumulating data on several designed sequences. We demonstrated that quadruplex-forming strands could be tailored to undergo a folding transition in a particular range of salt concentrations, and our results can be used to guide the design of nanoswitches for particular applications. For instance, if a quadruplex is required to switch at sub-mM concentrations of potassium, four quartets are needed. The choice of spacer is also important, as a poor choice would completely prevent the formation of quadruplexes.

Our investigation into the detection of triplexes using UV absorption spectroscopy revealed a drop in absorbance when triplexes are formed. This agrees with the findings of Brucale *et al* who also reported a drop in absorbance at 260 nm (40). The change is comparatively small,and would have been difficult to identify as a characteristic of triplex formation in the absence of fluorimetry data. It is important to note that we used only a single triplex-forming oligonucleotide, and the absorbance could vary based on sequence, triplex length, and other molecules in the solution. Although our triplex is comparatively short (10 nucleotides), 41% of all the bases are associated with the triple-stranded domain under triplex-forming conditions. Trying to detect triplex formation within larger sequences might be complicated by the other non-triplex nucleotides in the construct.

Determining triplex formation via UV spectra is not as conclusive as using fluorophore labelled oligonucleotides and fluorescence spectrophotometry, but UV absorbance spectroscopy is a label-free technique and the equipment required is less complex. Consequently, our findings may be of interest to researchers working with sequences that cannot be synthesised or labelled with fluorophores or those with more limited resources. We note that great care is needed with the UV absorbance measurements, with several repeats and careful cleaning needed.

## Conflict of interest

the authors declare no conflict of interest.

## Funding information

A.J.S. is funded by a PhD studentship from the University of Edinburgh. O.T.Y. is supported by an EPSRC iCASE award co-sponsored by DSTL. Additional funding was provided by an Institutional Strategic Support Fund (ISSF) grant from the Wellcome Trust.

## Author contribution statement

All authors contributed to development of methodology, performed analysis and contributed to preparation of the manuscript. BM – devised GQ sequences, performed original set of experiments on GQs, established experimental parameters. OTY – performed second set of experiments on GQs (data in manuscript), wrote scripts for data analysis. AJS – performed all experiments on the triplexes and associated controls. AE – supervisor to OTY. KED – conceived, supervised and managed this study.

## Notes

### Competing Interest Statement

The authors have declared no competing interest.

### Summary of Updates

Minor changes to figures (colour of lines in Fig. 4a, positioning of plots in Fig.5c). Clarification of Supplementary Datafile 1 - removal of unused dataset and inclusion of 'readme' sheet to explain how data is collapsed in other sheets.

