## Supplementary Information for "Using ultraviolet absorption spectroscopy to study nanoswitches based on non-canonical DNA structures"

**S1. G-quadruplex data**

All data is available in Supplementary Data File 1.

**S2. G-quadruplex data processing scripts**

Scripts are available at: https://github.com/OwenYeung/Non-canonical-DNA-structure

**S3. Triplex data**

All data is available in Supplementary Data File 2. All raw spectral data has been provided for each sample from both the nanodrop and fluorimeter in individual Excel sheets. Further Excel sheets have also been provided to give the averaged and standard deviation of each spectrum, and the peak data organized by pH (referred to as “processed”).

**S4. Data confirming linear range of Nanodrop**

Data is provided in Supplementary Data File 3.

**S5. Britton Robinson Buffer Compositions**

| **pH** | **0.4 M HCl (mL)** | **0.4 M CH_3_COOH**  **(mL)** | **0.4 M H_3_BO_3_ (mL)** | **0.4 M H_3_PO_4_ (mL)** | **0.04 M MgCl_2_ (mL)** | **0.2 M NaOH (mL)** | **0.4 M NaOH (mL)** | **UPW (mL)** |
| --- | --- | --- | --- | --- | --- | --- | --- | --- |
| **5** | 2 | 2 | 2 | 2 | 2 | 6.4 | - | 5.6 |
| **5.38** | 2 | 2 | 2 | 2 | 2 | 7.2 | - | 4.8 |
| **5.84** | 2 | 2 | 2 | 2 | 2 | 8 | - | 4 |
| **6.47** | 2 | 2 | 2 | 2 | 2 | 8.8 | - | 3.2 |
| **7** | 2 | 2 | 2 | 2 | 2 | 10 | - | 2 |
| **7.94** | 2 | 2 | 2 | 2 | 2 | - | 5.7 | 6.3 |
| **8.4** | 2 | 2 | 2 | 2 | 2 | 12 | - | - |
| **9** | 2 | 2 | 2 | 2 | 2 | - | 6.4 | 5.6 |
| **9.74** | 2 | 2 | 2 | 2 | 2 | - | 6.2 | 5.8 |
| **10.4** | 2 | 2 | 2 | 2 | 2 | - | 6.8 | 5.2 |
| **11.74** | 2 | 2 | 2 | 2 | 2 | - | 9 | 3 |

UPW = ultrapure water

**S6. Derivation of equation describing two-state folding kinetics**

NOTE: the folding process is believed to be much more complicated than this. Consequently, this should not be taken as an accurate representation of the underlying physics. The model derived is empirically useful and mathematically describes the phenomenon observed, but is a severe oversimplification (in part because it does not take into account the possibility of multiple folding topologies or kinetic traps).

Let us consider a two-state process, as follows:

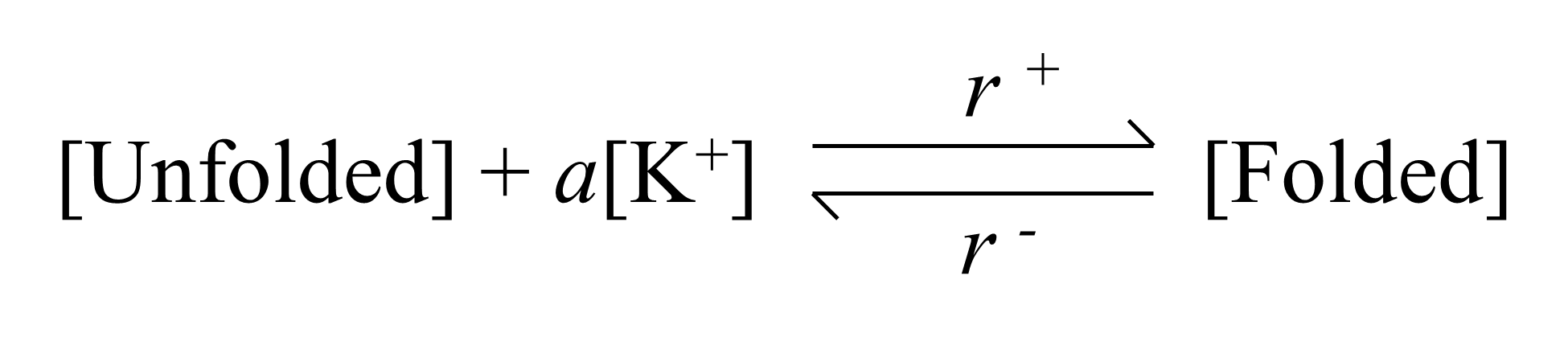

*a* represents the number of potassium ions that must be associated with the unfolded strand for folding to occur. The rate constants for the forward and backward reactions are represented by *r^+^* and *r ^–^* respectively, where we have avoided the usual notation of *k^+^* to avoid confusion with the potassium concentration.

The rate of change of the concentration of folded oligo is given by the following expression:

*d*[Folded]/*dt = r^+^*[Unfolded][K^+^]*^a^ – r ^–^* [Folded]

All our measurements are taken under equilibrium conditions, when the concentration of folded oligo does not change, and hence:

*r^+^*[Unfolded][K^+^]*^a^ = r ^–^* [Folded]

The total concentration of DNA is constant, which means that

[Unfolded] + [Folded] = [DNA]

Thus:

*r ^–^* [Folded] = *r^+^*([DNA] – [Folded] )[K^+^]*^a^*

Rearranging gives:

[Folded] = (*r^+^* [DNA] [K^+^]*^a^*) / (*r ^-^ + r^+^*[K^+^]*^a^*)

We assume that

*A_295_ = C* ([Folded]/[DNA]) + B ,

where C and B are constants.

Hence:

*A_295_*= (C*r^+^* [K^+^]*^a^*) / (*r ^-^ + r^+^*[K^+^]*^a^*) +B

When [K^+^] = 0, *A_295_* **=** *B*. Hence, we expect *B* ~ starting value for the dataset.

As [K^+^] tends to infinity, *A_295_* tends to *C+B*. Hence, *C* ~ ending value – starting value

Rearranging to cancel *r^+^,* we find:

*A_295_*= (C [K^+^]*^a^*) / (*E +* [K^+^]*^a^*) +*B*,

where *E* is the ratio of the rate constants. *E* is effectively the value of [K^+^] for which 50% of the oligos are folded.
